# Maternal separation disrupts noradrenergic control of adult coping behaviors

**DOI:** 10.1101/2025.03.09.642152

**Authors:** Chayla R. Vazquez, Léa J. Becker, Chao-Cheng Kuo, Solana A. Cariello, Ayah N. Hamdan, Ream Al-Hasani, Susan E. Maloney, Jordan G. McCall

**Author notes:** To whom correspondence should be addressed: J.G.M.: Address: 660 S. Euclid Ave, Box 8054, St. Louis, MO, USA, 63110 Website: www.mccall-lab.org.

## Abstract

Early life stress (ELS) in humans and preclinical rodent models profoundly impacts the brain and correlates with negative affective behaviors in adulthood. The locus coeruleus (LC), a stress-responsive brainstem nucleus that supplies most of the brain with norepinephrine (NE), is known to modulate negative affect. Here we used repeated maternal separation stress (MSS) to investigate the impact of ELS on the LC and stress-related behaviors in adulthood. Using *ex vivo* cell-attached electrophysiology, we recorded spontaneous LC firing across the lifespan from early development, pre-adolescence, adolescence, through adulthood. MSS significantly increased LC firing during early development and adulthood compared to No MSS mice. We next examined potential changes in the expression of genes linked to LC function. MSS decreased mRNA levels for both the alpha-2_A_ adrenergic receptor and dopamine beta-hydroxylase, the enzyme necessary for NE synthesis. At the behavioral level, MSS increased locomotion in approach-avoidance exploratory assays and increased immobility in the forced swim test. Forced swim increased LC cFos expression, a marker for neuronal excitation, in both No MSS and MSS mice. However, MSS mice had significantly less cFos than No MSS controls. We then sought to reverse this MSS-induced increase in immobility by inhibiting the LC during the forced swim test. In No MSS mice, LC inhibition increased immobility time, however, LC inhibition did not affect MSS immobility. Together, this study demonstrates that MSS dysregulates LC-NE activity across the lifespan and disrupts the role of the LC in regulating coping strategies during stressful events.

## Introduction

Early life stress (ELS) including neglect and emotional, physical, and/or sexual abuse is a major risk factor for poor mental and physical health throughout the lifespan [1–3]. Between 20 and 30% of adult psychiatric disorders can be attributed to ELS [4,5] and childhood maltreatment substantially increases the number of suicide attempts [6–9]. ELS is known to exacerbate stress-related behaviors and susceptibility to negative affective disorders specifically [9–11]. Childhood mistreatment has been shown to have a profound impact on stress-related brain regions such as the prefrontal cortex, hippocampus, hypothalamus, and the amygdala [8,12–14]. Furthermore, in rodent models, ELS leads to dysfunction in the hypothalamic-pituitary-adrenal axis and the central noradrenergic system, causing changes in stress reactivity [14,15]. In particular, recent studies have made it more clear that the locus coeruleus (LC) – norepinephrine (NE) system plays a critical role in early development and ELS [14,16–19].

The LC-NE system is a compact nucleus in the brainstem that extends widespread projections throughout the forebrain and spinal cord. It serves as the primary source of norepinephrine in the mammalian cortex and plays a key role in negative affective behaviors across species [20–30]. Indeed, acute or persistent types of stressors have long-term effects on LC activity and morphology which lead to increased negative affective behavior in adulthood [31,32]. ELS causes substantial, but poorly understood adaptation in the LC. For example, sleep deprivation in kittens decreased LC cell size and count [33]. Separately, re-exposure to stress in adulthood increases cFos, an immediate early gene associated with increased neuronal activity, in LC neurons of rats that went through maternal separation stress (MSS) [34]. Additionally, acute MSS changes spontaneous LC firing in rodents; with both increases [17] and decreases being reported [17,35]. These differences may be due to differences in the ELS model, species or developmental time chosen for recordings. Regardless, this suggests that ELS likely has a long-term impact on LC activity, but the mechanisms underlying these changes remain largely unknown.

How ELS affects LC activity is also potentially important for how long-lasting repercussions on behavior are sustained. In adult rodents, acutely enhanced LC activity has been shown to be necessary and sufficient for stressed-induced negative affective behaviors [26,29,36–40]. This increased spontaneous, tonic activity has also been described as suboptimal activity which correlates with suboptimal behavior [41,42]. This relationship between behavior and LC activity is often described as an inverted U-shaped curve. Optimal behavioral responses occur with moderate tonic LC activity, facilitating focused attention and cognitive flexibility. However, when LC activity is too low, insufficient norepinephrine release leads to inattentiveness (or suboptimal behavior). Conversely, too much LC activity also leads to suboptimal behavior such as heightened anxiety and distractibility. This relationship aligns with the Yerkes-Dodson law which states that performance improves only up to a point, after which too much arousal deteriorates performance [43,44]. Therefore, it is possible that ELS-induced negative affective behavior in adulthood could arise from either increases or decreases in LC activity. Investigating the long-term effects of ELS on the LC-NE system will thus give critical insights into the mechanisms underlying the behavioral outcomes of ELS.

Here we examine the effects of ELS, specifically MSS, on LC neurophysiology throughout the lifetime and adult negative affective behavior. We use *ex vivo* cell-attached electrophysiology and reverse transcription-quantitative polymerase chain reaction (RT-qPCR) to investigate the consequences of MSS on LC function. We find that MSS increases LC activity during early development and adulthood and decreases mRNA expression of important LC-NE genes. Behaviorally, MSS increases immobility in the forced swim test (FST), an assessment of coping behavior. To determine whether reducing this increased LC activity can change the effects of MSS-induced immobility, we used chemogenetics to inhibit the LC during FST. In mice that have had normal rearing and did not experience maternal separation (No MSS), LC inhibition increased immobility time. However, in MSS mice, the immobility time did not change with chemogenetic LC inhibition. Altogether, we show that MSS dysregulates the LC leading to altered coping behaviors in adulthood.

## Results

### Maternal separation increases locus coeruleus firing during early development and adulthood

Prior work in rats has shown that LC activity is altered 1-3 weeks after MSS combined with experimenter handling [17]. Importantly, however, the observed changes in LC activity varied depending on handling duration with short handling (15 minutes) decreasing LC spontaneous firing rate while longer handling (180 minutes) increased LC spontaneous firing rate [17]. Therefore, we sought to investigate the effects of MSS alone throughout the lifespan to determine when MSS-induced LC firing rate changes occur and how long these changes last. For our MSS paradigm, MSS mice underwent 4 hours of maternal separation from post-natal day (PND) 10 to PND 16, similar to previous work (except providing a normal amount of nesting material instead of limited amounts) [45,46]. We maintained No MSS mice in their normal breeding homecage environment, without experiencing MSS. We then performed *ex vivo* electrophysiological recordings from early development to adulthood (ages established through previous research [47]) following MSS (**Figure 1A-C**). In early development, MSS significantly increased LC firing rate (**Figure 1D**). This effect transiently disappeared during pre-adolescence and adolescence (**Figure 1E & F**) and was again observed in adulthood (**Figure 1G**). Together, MSS increased spontaneous LC activity shortly after MSS has occurred and a similar increase is reinstated in adulthood.

**Figure 1:**
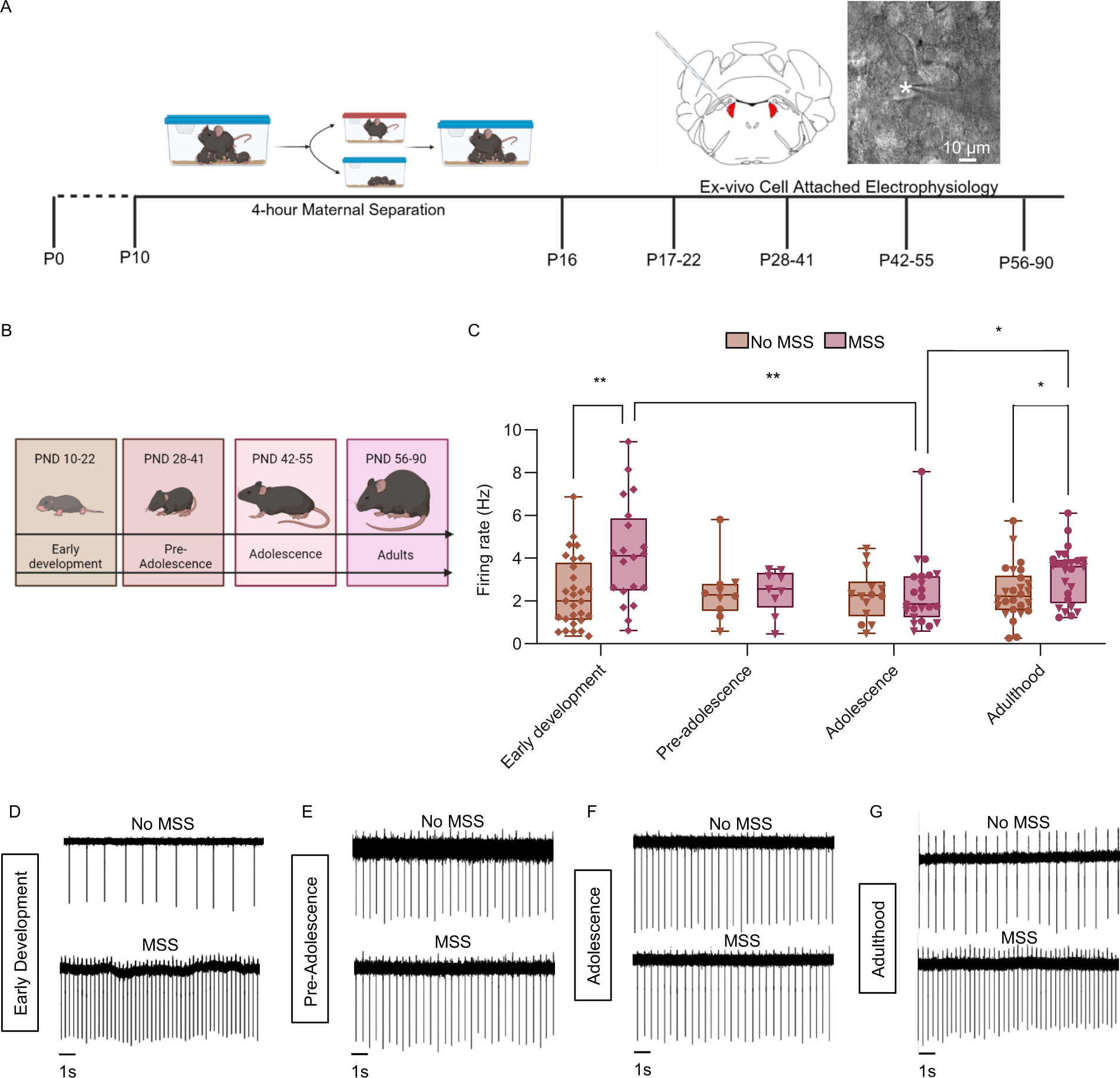
Maternal separation increases LC activity during early development and adulthood. (**A**) Timeline showing the maternal separation paradigm and electrophysiological time points, including a visual representation of the LC and electrode with a DIC image of a recorded LC neuron (*marks the pipette tip). (**B**) A diagram of mice at different developmental time points. (**C**) Firing rate of No MSS and MSS groups at all four life stages (Non-parametric analysis, Kruskal-Wallis for non-normal data with Mann-Whitney pairwise comparisons. KW: Group, X²(7,N=156)=18.874, p=.009 (Group variable has 8 levels, MSS and No MSS at each age)). Data are presented as a box and whiskers plot, where the box represents the interquartile range (IQR), the line within the box indicates the median, and whiskers extend from the minimum to maximum values. (**D-G**) Representative LC recordings from No MSS mice (top) and MSS mice (bottom) during early development (**D**), early adolescence (**E**), adolescence (**E**), and adulthood (**G**). Triangles ▼ in graphs indicate male animals; circles ● indicate female animals; ♦ indicate sex not recorded. Sex was not recorded for pups.

### Maternal separation decreases adult expression of genes associated LC-NE function

To determine what could be driving these electrophysiological adaptations seen in the LC after MSS, we examined genetic changes in the LC. We specifically focused on measuring gene expression for alpha-2_A_ adrenergic receptor (*Adra2a*), dopamine beta-hydroxylase (*Dbh*), tyrosine hydroxylase (*Th*), and corticotropin-releasing hormone receptor 1 (*Crhr1*). The activity of LC-NE neurons is in part regulated through alpha-2_A_ adrenergic receptors, which exert autoinhibitory feedback when norepinephrine is released [48,49]. Therefore, dysregulation of alpha-2_A_ adrenergic receptors could lead to changes in LC baseline activity. Additionally, prior work suggests that alpha-2_A_ receptor function is modulated after ELS in both human and animal models [50–52]. The two main enzymes in the NE biosynthesis pathway include TH and DBH [53]. TH the rate-limiting enzyme in the pathway converts tyrosine into L-DOPA, which is further converted into dopamine [54,55]. DBH acts downstream to convert dopamine into norepinephrine [55]. Both enzymes determine the availability of dopamine as a substrate for norepinephrine production, thus differential expression of these enzymes could give insight into LC function after maternal separation. Corticotropin-releasing hormone (CRH) is a key neuromodulator during a stress response [56,57]. CRH has also been shown to increase LC activity and CRH dysregulation has been associated with stress-related behaviors [58–60]. Thus, *Crhr1* expression could provide insight into whether the CRH system is being upregulated after MSS to increase LC activity.

Using RT-qPCR, we measured the expression of these genes in the adult LC region from MSS and No MSS mice (**Figure 2A&B**). MSS significantly decreased *Adra2a* and *Dbh* expression compared to No MSS mice. (**Figure 2C&D**). No significant differences, however, were found in *Th* or *Crhr1* expression between the two groups (**Figure 2E&F**). To identify potential downstream effects, we also examined adrenergic receptor expression in brain regions receiving inputs from the LC and relevant for stress-related behaviors including the basolateral amygdala, the central amygdala, and the anterior cingulate cortex [61–68]. Here we saw no changes in downstream adrenergic receptor mRNA expression (**Figure S1**). Together it appears that decreased *Adra2a* expression in the LC could contribute to the increase in LC firing that we observed in adulthood after MSS.

**Figure 2:**
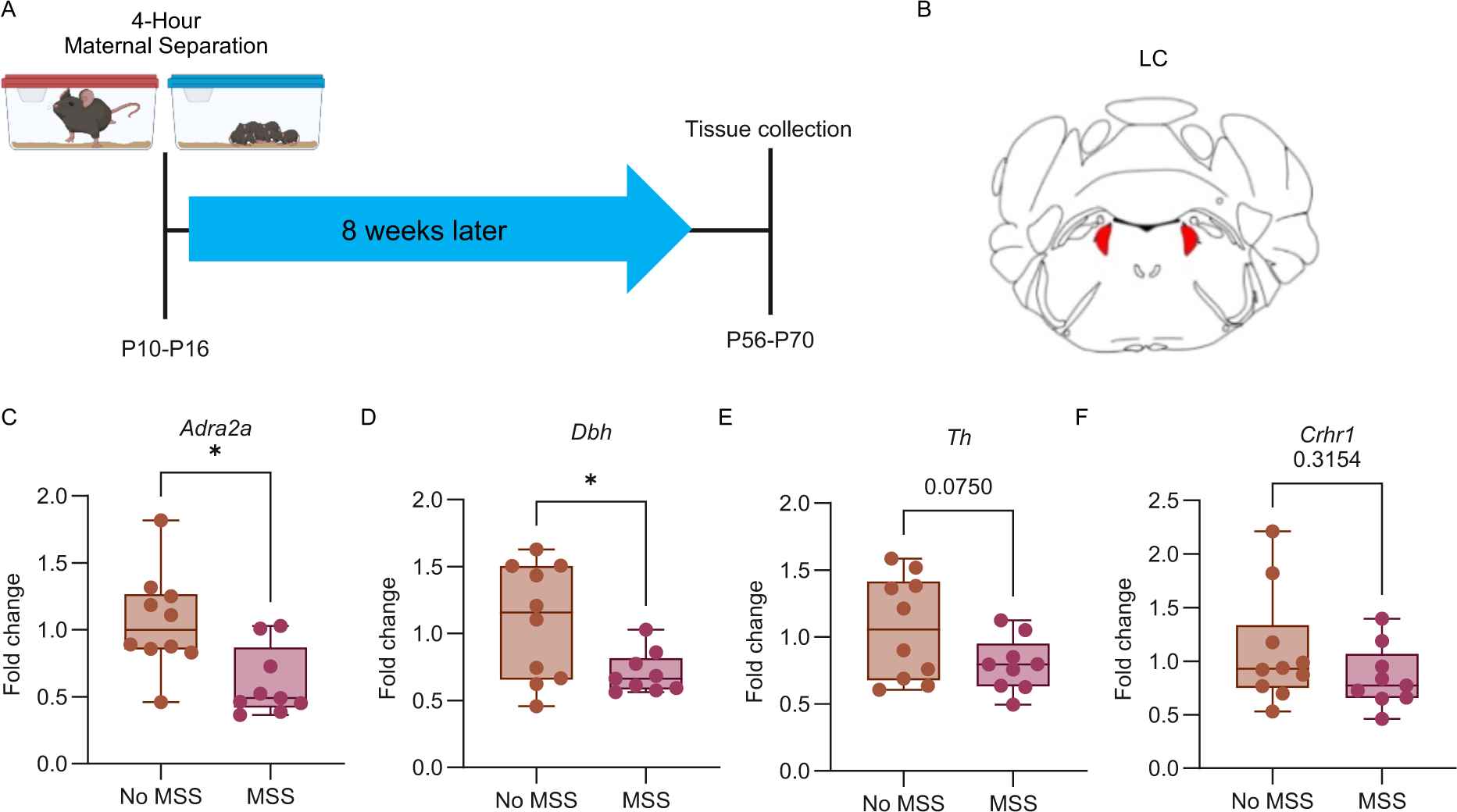
MSS alters mRNA content of the LC. (**A**) Timeline showing the time of tissue collection for the mRNA extraction after MSS. No MSS mice also had their tissue collected within the same age range. (**B**) Schematic of LC tissue collection. (**C**) *Adra2a* expression in the LC was significantly decreased in MSS animals compared to No MSS animals (Mann-Whitney test, U=14, p=0.0101). (**D**) *Dbh* expression in the LC was significantly decreased in the MSS animals compared to No MSS animals (Student’s t-test, t(17) = 2.503, p=0.0228). There was no difference between in *Th* (Student’s t-test, t(17) = 1.897, p=0.0750) (**E**) or *Crhr1* (Mann-Whitney test, U=32, p=0.3154) (**F**) expression in the LC between MSS and No MSS mice. Triangles ▼ in graphs indicate male animals; circles ● indicate female animals. Data are presented as box and whiskers plots, where the box represents the IQR, the line within the box indicates the median, and whiskers extend from the minimum to maximum values.

### Maternal separation increases adult passive coping in the forced swim test

After identifying increased LC spontaneous firing rate and downregulation of *Adra2a* and *Dbh* expression in the LC in adult mice after MSS, we sought to determine whether these changes influenced negative affective behaviors associated with LC function. We subjected MSS and No MSS mice to three different types of stress-related behavior tests: the elevated plus maze (EPM), open field test (OFT), and forced swim test (FST) (**Figure 3A**). The EPM tests anxiety-related behavior by exploiting the approach-avoidance conflict. Mice who spend more time in the open arms are said to show more approach behavior compared to mice who spend more time in the closed arms which show more avoidance behavior. Additionally, acutely increased LC activity produces avoidance behavioral responses in the EPM and the similar elevated zero maze [27–30,36,58,69]. Furthermore, using antisense oligonucleotides to reduce *Adra2a* expression in the LC increases open-arm entries in the EPM in male rats compared to No MSS mice [70]. Like the EPM, the OFT also examines anxiety-related behavior, and observations within this test have been linked to LC function [26,27,71,72]. The OFT is also well-suited to observe changes in locomotor activity caused by LC dysregulation. Finally, the FST measures changes in coping behavior during a stress-inducing experience. Animals that are more mobile during the test are considered to show active coping behavior while those that spend more time immobile are said to show passive coping behavior [73–77]. These behaviors during FST are modulated by noradrenergic activity [39], specifically one study showing that activating alpha2a decreases immobility [78]. Together these three behavioral measures may provide some insight into LC function following MSS.

**Figure 3:**
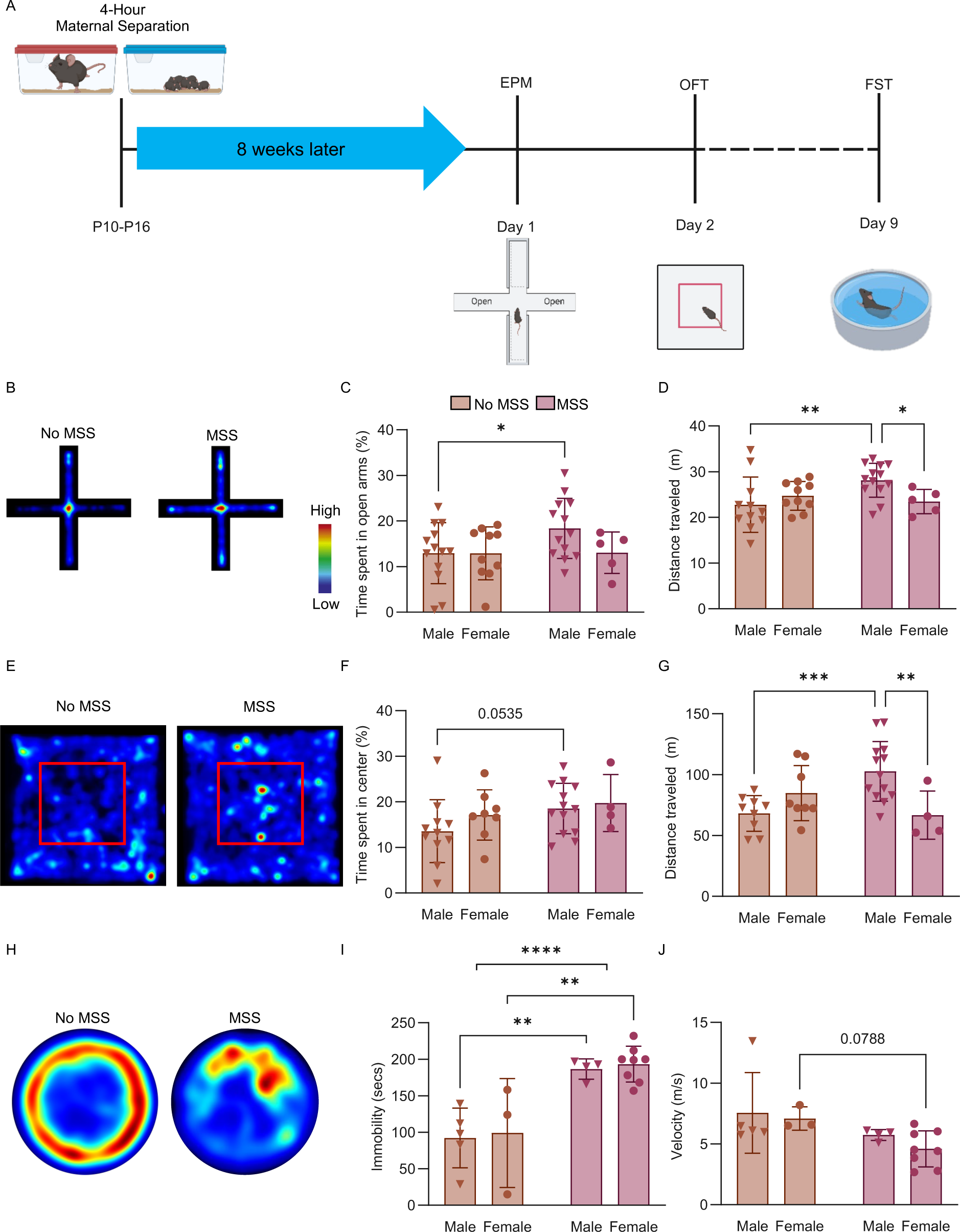
Maternal separation increases passive coping behavior during Forced Swim Test. (**A**) Timeline showing the timing of behavioral tests 8 weeks after maternal separation. (**B**) Representative heat maps showing activity during the elevated plus maze. (**C**) The percent of time spent in the open arms of the elevated plus maze (Two-way ANOVA, F(1, 37) = 1.739, p= 0.0337). (**D**) The total amount of distance moved in the elevated plus maze (Two-way ANOVA, F(1, 35) = 5.048, **p= 0.0045 and *p= 0.047). (**E**) Representative heat maps showing activity during the open field test. (**F**) The percent of time spent in the center of the open field test (Two-way ANOVA, F(1, 32) = 2.881). (**G**) The total distance moved in the open field test (Two-way ANOVA, F(1, 30) = 10.78, ***p=0.0008 and **p= 0.0063). (**H**) Representative heat maps showing activity during the forced swim test. (**I**) The amount of time spent immobile during the forced swim test for the No MSS and MSS group (Two-way ANOVA, F(1, 16) = 27.70,****p= <0.0001, **p= 0.0018 and **p=0.0019, respectively. (**J)** The average velocity during the forced swim test for the No MSS and MSS group (Two-way ANOVA, F(1, 16) = 5.314). Triangles ▼ in graphs indicate male animals; circles ● indicate female animals. All data represented as mean ± SEM.

In the EPM, MSS mice spent more time in the open arms compared to No MSS mice (**Figure 3B&C**; **Figure S2A**). This effect appears to be largely driven by male MSS mice (**Figure 3C**). Additionally, male MSS mice had increased locomotion compared to both female MSS and No MSS males (**Figure 3D; Figure S2B**). Other recent studies have also shown that maternal separation causes an increase in time spent in open arms [79,80]. We found no changes in time spent in the center of the OFT between MSS and No MSS animals (**Figure 3E&F**; **Figure S2C**). Furthermore, we again observed that MSS males displayed an increase in total distance traveled compared to non-MSS males in OFT (**Figure 3G; Figure S2D**). Finally, in the FST, MSS significantly increased immobility time in both sexes without significantly altering velocity (**Figure 3H-J**), indicating MSS increases passive coping behavior in mice. From these tests, the FST was the only assay where both female and male MSS mice significantly differed from No MSS mice, thus making it the most selective indicator for MSS-altered, potentially LC-related negative affective behavior.

### Forced swim test activates LC neurons regardless of prior maternal separation

Given that MSS most prominently affected performance in the FST, we aimed to understand whether the LC is differentially activated in this test across groups. To do so, we measured the expression of cFos, a marker of neuronal activation in the LC [81], in both MSS and No MSS animals immediately following FST compared to mice that did not go through the FST (**Figure 4A**). Using immunohistochemistry to stain for cFos and positive TH cells in the LC, we counted cFos within an anatomically defined ROI that encapsulated the LC (**Figure S3**). cFos significantly increased in both No MSS and MSS groups that went through FST compared to non-FST No MSS mice (**Figure 4B&C**). Interestingly, MSS mice show significantly fewer LC cFos+ cells within the LC after FST compared to No MSS. Thus, these results show that LC is recruited and active during FST, but that MSS might alter LC response to forced swim.

**Figure 4:**
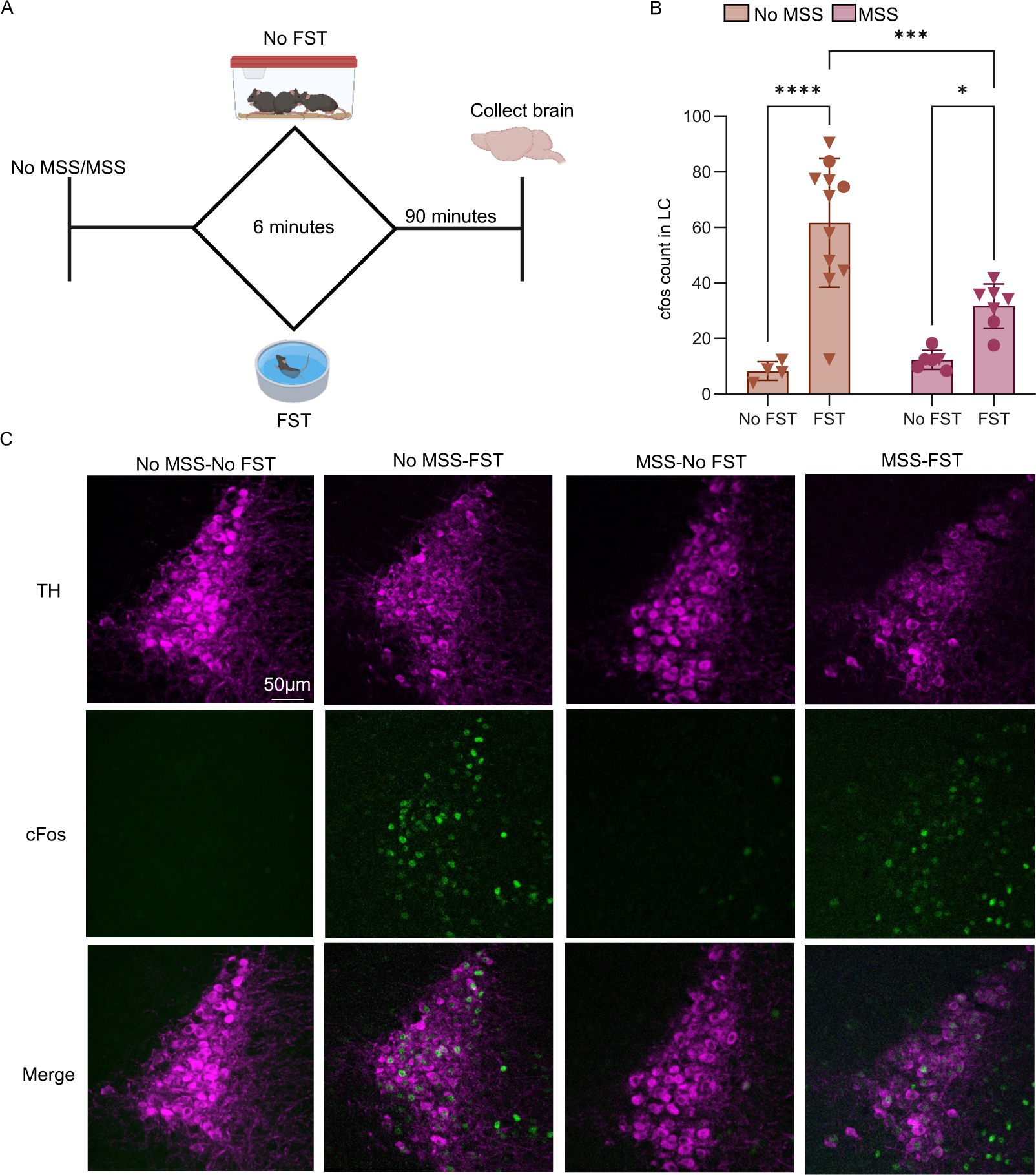
Forced Swim Test increases LC cFos expression. (**A**) Timeline for quantification of cFos expression within LC after force swim test. The FST mice went through FST for 6 minutes and were sacrificed 90 minutes afterward. The No MSS mice did not go through FST and instead sat outside of their colony room for 6 minutes. After 90 minutes they were sacrificed. (**B**) The cFos count in the LC per animal within each group (Data represented as mean ± SEM, Two-way ANOVA test, F(1, 24) = 7.287, ****p<0.0001, ***p= 0.0006, * p=0.0348). (**C**) Confocal images of the cFos. Triangles ▼ in graphs indicate male animals; circles ● indicate female animals.

### MSS prevents LC inhibition-mediated passive coping behavior

After observing that the LC is recruited during FST and knowing that LC baseline firing rate is significantly increased after MSS, we sought to reverse the effects of MSS-induced immobility during FST by inhibiting the LC. To do so we injected a virus to selectively express either inhibitory DREADD receptors (designer receptors exclusively activated by designer drugs; hM4Di; AAV8-hSyn-DIO-hM4D(Gi)-mCherry) or mCherry alone (AAV8-hSyn-DIO-mCherry) in the LC of *Dbh*^Cre^ mice (**Figure 5A**). We used whole-cell current clamp to functionally validate hM4Di expression (**Figure 5B-H**). Here, bath application of CNO significantly increased the amount of current needed to elicit an action potential (i.e., the rheobase) in hM4Di+ LC neurons, but not in mCherry+ LC neurons (**Figure 5C-F**). Furthermore, in response to increasing current, the firing rate for mCherry did not change with CNO application (**Figure 5G**). However, CNO application on hM4Di-expressing neurons caused a significant rightward shift in firing rate (**Figure 5H**). Together, this increased rheobase and decreased firing in response to current suggests that CNO-mediated activation of hM4Di decreases LC excitability. We next sought to test whether blunting LC excitability in MSS animals could rescue the MSS-induced increased immobility during FST (**Figure 5I&J**). To do so, we administered CNO (3 mg/kg, i.p.) or saline 30 minutes prior to FST in both No MSS and MSS mice expressing hM4Di (**Figure 5I)**. In No MSS mice, CNO increased immobility. Interestingly, however, CNO had no effect on MSS mice (**Figure 5J**). This suggests that MSS possibly hinders the LC’s ability to regulate passive coping strategies.

**Figure 5:**
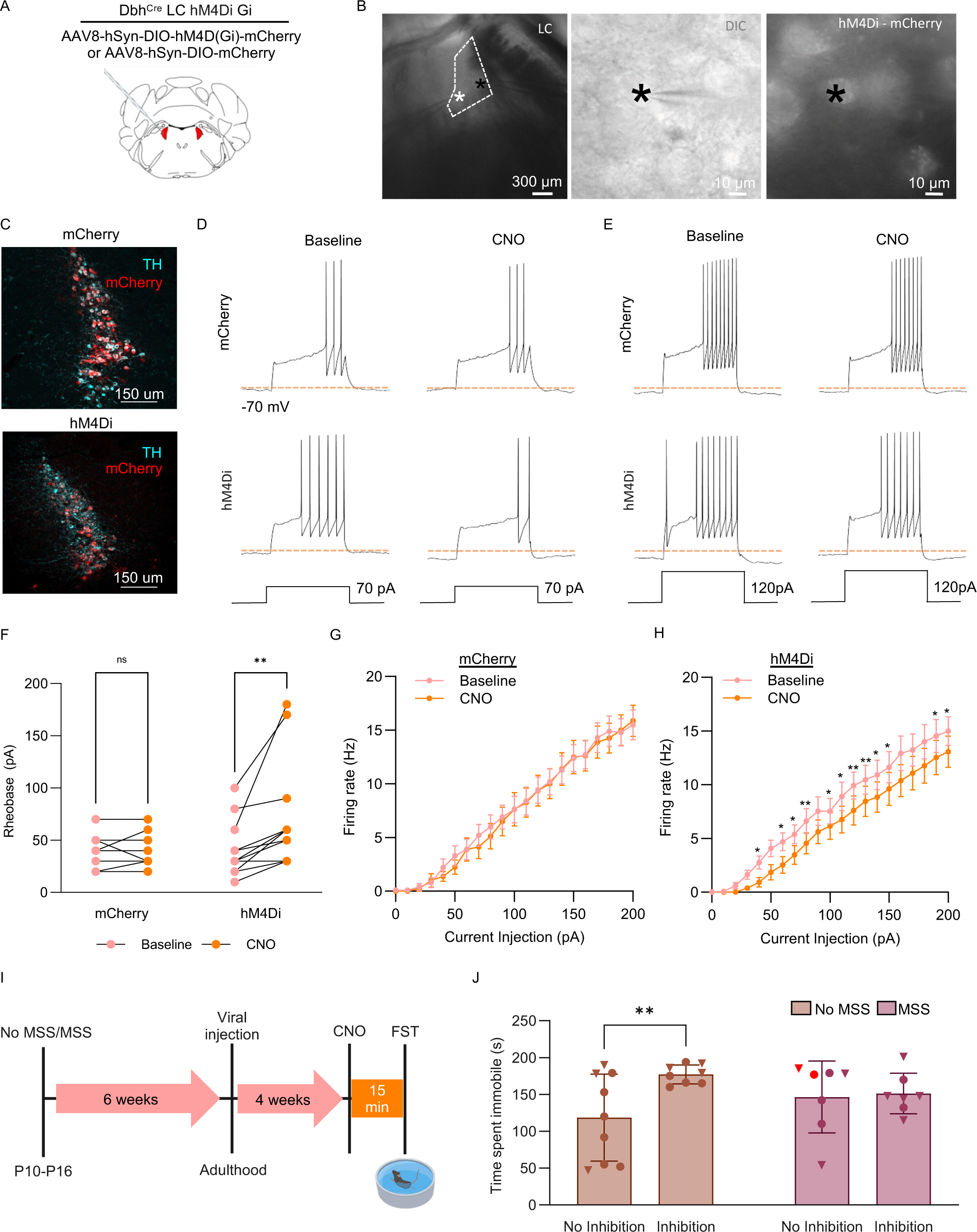
Inhibiting the LC in MSS mice during FST rescues MSS-driven passive coping behavior. (**A**) A diagram of the LC where the bilateral injection occurs. (**B**) Left, a DIC image under low magnification of the LC with the two recording pipettes marked by asterisks. Middle, an image for one of the recorded cells shown in left panel indicated by a black asterisk. Right, a fluorescent image of the same recorded LC neuron showing hM4Di-mCherry expression. (**C**) Confocal image showing the colocalization of mCherry or hM4Di-mCherry with *TH* immunoreactive signals. (**D-E**) Representative traces from a mCherry and hM4Di-mCherry expressing LC neurons at baseline and with CNO administration upon current injection at 60 (**D**) and 100 (**E**) pA. (**F**) Graph showing the rheobase in mCherry and hM4Di-mCherry expressing LC neurons at baseline and under CNO administration (Two-way ANOVA, F(1, 27) = 4.343, **p = 0.0013). (**G-H**) Plots demonstrating input-output relationship from current injections in mCherry (G) and hM4Di-mCherry (H) expressing LC neurons at baseline and after CNO is administered (G: Two-way ANOVA, F (1, 16) = 0.03017; H: Two-way ANOVA, F(1, 24) = 1.580, *p<0.05 and **p<0.01). (**I**) An experimental timeline for the behavioral examinations. (**J**) A graph showing the immobility in force swim test from No MSS and MSS animals, all animals expressing hM4Di-mCherry received either saline or CNO prior to the test except two animals MSS no inhibition animals that express mCherry and received CNO (denoted in red).

## Discussion

Here we identified that repeated maternal separation leads to increased LC activity during early development and adulthood. Interestingly, this increased activity transiently reversed during pre-adolescence and adolescence suggesting potentially different mechanisms underlying the increased activity in early development and adulthood. In adulthood, MSS decreased *Dbh* and *Adra2a* mRNA levels in the LC. We next assessed negative affective behaviors in adulthood and showed that MSS increased locomotion in male mice and increased passive coping behavior in both sexes. Lastly, while LC inhibition in No MSS mice increased passive coping, LC inhibition in MSS mice did not alter behavior in the FST.

The literature on stress-related and coping behaviors after MSS is inconsistent. One strong possibility for the discrepancies across studies is the developmental timepoint(s) at which MSS occurred. It is not yet clear how these differences could impact LC activity and behavior. For instance, one study administered MSS from PND 2 to PND 12 in a strain of C57BL/6 mice, earlier than our manipulation from P10-P16 and found no effects in time spent in the open arms of the EPM or the time spent in the center area of the OFT during adolescence or adulthood [82]. Despite these contrasts with our findings, this study did identify significantly increased immobility during the FST during adolescence. While we did not test behavior during adolescence, we did observe this effect in adulthood. In another study, MSS was administered from PND 2 to PND 14 in a strain of C57BL/6 mice, again earlier than our MSS, resulting in decreased open arm time in the EPM for MSS females compared to No MSS mice [83]. Additionally, this study showed MSS males decreased locomotor activity in the OFT compared to No MSS mice – the opposite effect we observed here. Other work showed that C57BL/6J pups that experienced MSS during the late postnatal period (PND 10 to PND 20, running later than our MSS) and then experienced social defeat had decreased social interaction, decreased sucrose preference, and increased immobility during FST compared to pups that had MSS during the early postnatal period (PND 2 to PND 12) which were proportionate to No MSS mice [45]. More recently, a similar MSS timeline (PND6 to PND 16) combined with early weaning and adolescent stress showed increased locomotion during EPM, but decreased time spent in open arms in female MSS BALB/cJ mice [35]. Remarkably, in contrast to our findings, this study showed decreased LC excitability following these early and adolescent stressors. Although our data in C57BL/6J mice align with earlier research showing an increase in LC activity after MSS [17], a follow-up study from the same group using C57BL/6J mice showed similar effects as well as differences in males and females possibly arising from sexual dimorphic responses to CRH [84,85]. Together these apparently conflicting reports suggest that LC adaptation to ELS is dependent on the timepoint and duration of MSS, the length of each separation, and the inclusion of other stressors. Furthermore, ultimate understanding is likely more complex than a binary increase or decrease in tonic firing, which is consistent with our evolving understanding of LC firing across strains and species [86].

To try to understand why LC activity is increased in our adult MSS animals, we performed RT-qPCR for several LC-related genes. Many studies show differential gene expression in other brain regions following ELS [45,87–90]. For example, animals that went through MSS from PND 5 to PND 21 for 6 hours each day appear to have a higher level of *Crh* in the hypothalamus compared to No MSS mice [91]. This increased hypothalamic *Crh* could have functional consequences for the LC and its associated circuitry. In our analysis, however, *Crhr1* mRNA was not changed. While this does not rule out the possibility of receptor desensitization or other translational and post-translational modifications in the CRH system, MSS did decrease *Adra2a* mRNA in LC. Loss of alpha-2_A_-mediated autoinhibition could contribute to increased LC activity following MSS. The decrease in *Dbh* could reflect a compensatory downregulation of enzymatic activity or the possibility that adrenergic signaling in downstream regions could be altered, we did not, however, observe any significant changes to the adrenergic receptor genes we probed in downstream regions (**Supplementary Figure 1**). Sleep deprivation in kittens reduces LC cell counts [33]. It is therefore possible that the MSS-driven decreases in *Adra2a* and *Dbh* could be driven by few LC neurons, however, in such a case we would also expect a decrease in *Th* and *Crhr1* which we did not observe. It is also possible that the decrease in *Dbh* could lead to increased levels of dopamine since *Dbh* is necessary to metabolize dopamine into norepinephrine. To fully understand the implications of these changes, further research would be necessary to assess functional changes in NE release and receptor activity at different time points such as directly after MSS, in early development. Future efforts should also examine LC genetic changes in more depth for other possible mechanisms that could contribute to the MSS-induced dysfunctional LC activity.

The central noradrenergic system is known to modulate coping behaviors in FST. For example, infusion of norepinephrine into the LC of naïve rats had a non-linear effect on immobility with some doses increasing and other doses decreasing immobility in the FST [38]. Subsequent work demonstrated that alpha2a-mediated immobility during forced swim is driven by alpha2a receptors on non-LC cells [78]. However, another study showed inhibiting TH by a-methyl-para-tyrosine methyl ester (AMPT) increases immobility for mice that have transgenic, lifelong corticotropin-releasing hormone (CRH)-mediated hyper-activation of the locus coeruleus [59]. These latter results appear to align with our study showing chronically enhanced LC activity and downregulated *Dbh* expression. To determine the functional relevance of this increased activity, we sought to inhibit the LC in MSS animals that have increased LC firing. Inhibiting the LC in No MSS animals increased immobility similar in scale to how MSS increased immobility. While these results appear at odds (i.e., hM4Di decreases LC excitability and MSS increases LC firing), both phenomena push the LC activity outside of the optimal firing rate range into hypo- and hyper-firing rate ranges, respectively. This appears to be a possible example of the Yerkes-Dodson law where both extremes lead to dysfunctional behavior [92,93]. Therefore, we tested to see if inhibiting the LC in MSS mice would bring the LC closer to normal firing rates and thus reverse the suboptimal behavior we see in FST potentially caused by the MSS-induced increases in LC activity. However, we did not see a decrease in immobility time, which possibly supports the hypothesis that pushing the LC activity into a suboptimal tonic firing rate range can inhibit the ability of the LC to modulate behavior normally. It is also possible that decreased excitability in slice is overwhelmed by endogenous synaptic input *in vivo* in MSS mice, rendering the chemogenetic manipulation less functional. Many studies have investigated the effects of manipulating LC tonic and phasic activation on different behaviors through optogenetic activation [26,94–96]. However, not many have tested manipulating LC activity after it is naturally firing at a suboptimal rate to see changes in behavior. Altogether, this study provides insight into how dysregulated LC activity after MSS could prevent the LC from optimally controlling behavior.

## Methods

### Animals

Male and female C57BL/6J (JAX:000664) and Dbh-Cre (JAX:033951) mice were used. Mice were initially sourced from The Jackson Laboratory (Bar Harbor, ME, USA) and bred in-house in a barrier facility in another building. Adult animals were either transferred to a holding facility adjacent to the behavioral space between 4-6 weeks of age or bred directly in the holding facility. Pups used in the experiments were only bred in the holding facility. All mice were group-housed, given *ad libitum* access to standard laboratory chow (PicoLab Rodent Diet 20, LabDiet, St. Louis, MO, USA) and water, and maintained on a 12:12-hour light/dark cycle (lights on at 7:00 AM). All experiments and procedures were approved by the Institutional Animal Care and Use Committee of Washington University School of Medicine in accordance with National Institutes of Health guidelines.

### Maternal Separation Stress

The dam is placed in a new separate cage with regular bedding, food, and water for 4 hours during a one-week period in a separate room. The pups are left in their home cage on a heating table (∼42°C) to avoid hypothermia. This week starts on postnatal day 10 and ends after postnatal day 16. The time of day the maternal separation takes place is purposely varied throughout the week to avoid habituation to the procedure. The sire is separated from the cage before the maternal separation week starts.

### Acute Slice Preparation

Adult mice were deeply anesthetized via a i.p. injection of a mix of ketamine (69.57 mg/ml), xylazine (4.35 mg/ml), & acepromazine (0.87 mg/ml). Upon sedation, adult mice were perfused with slicing-aCSF consisting of 92 mM N-methyl-d-glucose (NMDG), 2.5 mM KCl, 1.25 mM NaH_2_PO_4_, 10 mM MgSO_4_, 20 mM HEPES, 30 mM NaHCO_3_, 25 mM glucose, 0.5 mM CaCl_2_, 5 mM sodium ascorbate and 3 mM sodium pyruvate, oxygenated with 95% O_2_ and 5% CO_2_. pH of aCSF solution was 7.3–7.4. Pups were anesthetized via exposure to isoflurane that was dropped on tissue in a mouse transfer container. Upon sedation, pups were decapitated, and the cranium was submerged in aCSF immediately after. In both adults and pups, the brain was dissected and embedded with 2% agarose in slice-aCSF. Coronal brain slices were cut into 300 μm slices using a vibratome (VF310-0Z, Precisionary Instruments, MA, USA) and incubated in warm (32°C) slicing-aCSF for 10 mins. After incubation slices were transferred to holding-aCSF containing 92 mM NaCl, 2.5 mM KCl, 1.25 mM NaH_2_PO4, 30 mM NaHCO_3_, 20 mM HEPES, 25 mM glucose, 2 mM MgSO_4_, 2 mM CaCl_2_, 5 mM sodium ascorbate and 3 mM sodium pyruvate, oxygenated with 95% O_2_ and 5% CO_2_ for one hour. pH of the solution was 7.3–7.4. Slices were placed into a recording chamber mounted on an upright microscope (BX51WI, Olympus Optical Co., Ltd, Tokyo, Japan) with epifluorescence equipment and a highspeed camera (ORCA-Flash4.0LT, Hamamatsu Photonics, Shizuoka, Japan) while perfused continuously with warm (29–31°C) recording-aCSF containing 124 mM NaCl, 2.5 mM KCl, 1.25 mM NaH_2_PO_4_, 24 mM NaHCO_3_, 5 mM HEPES, 12.5 mM glucose, 2 mM MgCl_2_, 2 mM CaCl_2_, oxygenated with 95% O_2_ and 5% CO_2_ and pH 7.3–7.4.

### Electrophysiology

All recordings were performed using visual guidance (40× water immersion objective lens, LUMPLFLN-40xW, Olympus, Tokyo, Japan) through a glass pipette pulled from borosilicate glass capillary (GC150F-10, Warner Instruments, Hamden, CT, USA) with a resistance from 2-5 MΩ. All data were collected using a Multiclamp 700B amplifier (Molecular Devices, San Jose, CA, USA) with a low-pass filtered at 2 kHz and digitized at 10k Hz through Axon Digidata 1440A interface (Molecular Devices, CA, USA) running Clampex software (Molecular Devices, CA, USA). Electrophysiology data was exported through Clampex software and analyzed using Easy Electrophysiology (Easy Electrophysiology Ltd, London, England) and GraphPad Prism 10 (GraphPad Software, MA, USA). <u>Cell-attached recordings</u>: Cells were recorded using the cell-attached recording method in voltage-clamp mode with pipettes filled with recording-aCSF. <u>Whole-cell recordings</u>: For the hM4Di validation experiments, glass pipettes were filled with potassium gluconate-based intra-pipette solution consisting of 120 mM potassium gluconate, 5 mM NaCl, 10 mM HEPES, 1.1 mM EGTA, 15 mM Phosphocreatine, 2 mM ATP and 0.3 mM GTP, pH 7.2–7.3 and osmolality adjusted to 300 mOsm. A green LED light was used to identify viral expression in neurons before recording, targeted LC neurons were recorded under current clamp mode. For examinations of input-output relationship, current injections from -50 to 200pA with 10pA steps, were applied while membrane potential was controlled between -70 to -75mV. CNO was delivered through the recording-aCSF perfusion system.

### Elevated Plus Maze (EPM)

Mice were habituated in the room where the experiment was taking place for 30 minutes before the start of the experiment. The EPM is a plus sign-shaped platform elevated off the ground with four arms: two enclosed with walls and two open with no walls attached. The experiment ran for 11 minutes in dim lighting (8-10 lux). The anxiety-related behavior is measured by the sum amount of time spent in the open arms. The maze was cleaned with 70% ethanol between each trial.

### Open Field Test (OFT)

Mice were habituated in the room where the experiment was taking place for 30 minutes before the start of the experiment. OFT testing was performed in a 50 x 50 cm square enclosure for 20 minutes in dim lighting (8-10 lux); the center is defined as a square that is 50% of the total OFT area. The open field was cleaned with 70% ethanol between each trial.

### Forced Swim Test (FST)

Mice were placed in an opaque 5-liter beaker (30 cm tall x 18 cm in diameter) filled with 3.5 L of 26 ± 1 °C water. Animals were placed in the water to swim for a single, 6-minute trial. After, the mice were paper towel dried and placed on a covered heating pad to prevent hypothermia, then returned to their home cage. The last 4 minutes of the trial were analyzed for time spent immobile.

### Immunohistochemistry Staining for cFos

Mice were anesthetized with an i.p. injection of a cocktail containing ketamine (69.57 mg/ml), xylazine (4.35 mg/ml), & acepromazine (0.87 mg/ml). They were then perfused with 20 ml of cold 1 x PBS and then 20 ml of cold 4% paraformaldehyde in 0.1M PB. Brains were dissected and postfixed in paraformaldehyde for 24 hours at 4°C. After, the brains were left in 30% sucrose in 0.05M PB for 72 hours. Next, the brains were frozen and sliced into 30 μm thick sections using a microtome (SM2000R, Leica, Germany). Sections were rinsed three times with PBS and then incubated in a blocking solution containing 2% bovine serum albumin (BSA) plus 5% normal goat serum (NGS) in PBST for one hour before transferring to a PBS solution containing c-Fos (9F6) (rabbit, 1:1000, Cell Signaling Technology, 2250) and tyrosine hydroxylase (TH) (#TYH, 1:1000; Aves Labs Inc., Tigard, OR, USA) primary antibodies overnight on a shaker at room temperature. Sections were then washed in PBS 3 times and incubated in PBS with secondary antibodies Alexa Flour 488 anti-rabbit (1:400; Cat#A-11008; Invitrogen, Carlsbad, California, USA) and Alexa Flour 594 anti-chicken (1:1000; Cat#A-11042; Invitrogen, Carlsbad, California, USA) for 2 hours at room temperature followed by a rinse with PBS for three times. Sections were then mounted on glass slides with Vectashield mounting medium (Vector Labs, CA, USA). Images were collected using a Leica confocal microscope (SP8, Leica, Germany).

### Viral Preparation

pAAV-hSyn-DIO-hM4D(Gi)-mCherry was a gift from Bryan Roth (Addgene viral prep # 44362-AAV8). pAAV-hSyn-DIO-mCherry was a gift from Bryan Roth (Addgene viral prep # 50459-AAV8) [97].

### Stereotaxic Surgery

Mice were anesthetized in an induction chamber (3% isoflurane) and placed in a stereotaxic frame (Kopf Instruments, Model 940) where they were maintained at 2-2.5% isoflurane throughout the procedure. A craniotomy was performed, and mice were injected with 350 nL of AAV8-hSyn-DIO-hM4D(Gi)-mCherry or AAV8-hSyn-DIO-mCherry bilaterally into the locus coeruleus. Stereotaxic coordinates from bregma: -5.45 mm anterior-posterior (AP), ± 1.10 mm medial-lateral (ML), and -3.75 mm dorsal-ventral (DV) were used. Postoperative care included access to ¼ carprofen tablet (5 mg/kg) per mouse and a 0.15 mL subcutaneous saline injection immediately following surgery. To allow for sufficient viral expression, mice were allowed to recover for at least four weeks prior to behavioral testing and slice electrophysiology experiments.

### Tissue collection for qPCR

#### Locus coeruleus

Mice were anesthetized with a mix (i.p. 182 mg/kg) of ketamine (69.57 mg/ml), xylazine (4.35 mg/ml), & acepromazine (0.87 mg/ml) and perfused with ice-cold aCSF containing the following (in mM): 92 N-methyl-d-glucose (NMDG), 2.5 KCl, 1.25 NaH_2_PO_4_, 10 MgSO_4_, 20 HEPES, 30 NaHCO_3_, 25 glucose, 0.5 CaCl_2_, 5 sodium ascorbate and 3 sodium pyruvate, oxygenated with 95% O_2_ and 5% CO_2_. pH of aCSF solution was 7.3–7.4 and osmolality adjusted to 315–320 mOsm with sucrose. The brainstem was dissected and embedded with 2% agarose in aCSF and coronal brain slices were cut into 250 μm slices using a vibratome (VF310-0Z, Precisionary Instruments, MA, USA). Bilateral LC was dissected under a microscope (Leica S6E, Leica Microsystem GmbH), immediately frozen, and kept at -80 °C until RNA extraction.

#### Other tissues

Following brainstem isolation, the remaining rostral part of the brain was cut into 1mm thick coronal slices using a brain matrix (Zivic Instruments). Slices containing the ACC, CeA, and BLA were transferred in a petri dish filled with aCSF and put under a microscope. Bilateral ACC were dissected using anatomical landmarks. Bilateral CeA and BLA were punched using 20g blunt syringe needle (Warner Instruments). All tissues were immediately frozen on dry ice and kept at -80 °C until RNA extraction.

#### Locus coeruleus, central amygdala and basolateral amygdala

Total mRNA was extracted from tissue using the Arcturus PicoPure RNA Isolation Kit (Thermo Fisher Scientific, Waltham, MA). Around 1 mg of LC tissue was incubated for 30 min in 50 µl of extraction buffer at 42 °C, (500 rpm) and then centrifuged for 2 min at 3,000 g. The supernatant (50 µl) was transferred in a new tube containing 50 µl of 70% EtOH. The mix was then loaded on an RNA Purification Column (pre-conditioned with 250 µl of conditioning buffer for 5 min), centrifuged at 100 g for 2 min (binding of the RNA to the column), and at 16,000 g for 30 s to remove the flowthrough. Next 100 µl of washing buffer 1 (WB1) was added and centrifuged at 8,000 g, for 1min, before adding 10 µl of DNAse and 30 µl of RDD buffer (Qiagen, Germany). The mix was left at room temperature for 15 min before adding 40 µl of WB1 and centrifuge at 8,000 g for 15 s. The column was then washed two times by adding 100 µl of washing buffer 2, centrifuged at 8.000 g for 1 min after the first wash, and two times at 16.000 g for 1 min after the second wash to remove all traces of buffer. Columns were transferred to a new 0.5 ml collection tube, to which 12 µl of elution buffer was added, left at room temperature for 1 min, and centrifuged at 1.000 g for 1 min to distribute the elution buffer on the column. Finally, the RNA was eluted by centrifugation for 1 min at 16.000 g. RNA concentrations were measured by spectrophotometry (Nanodrop One, Thermo Fisher Scientific, Waltham, MA). Samples were then kept at −80 °C until use.

#### Anterior cingulate cortex

Total mRNA was extracted from tissue using the Qiagen RNeasy minikit (Qiagen, Germany). Around 20mg of tissue were transferred to bead lysis 1.5 mL tubes (Pink RINO RNA lysis kit, Next Advance, Troy, NY) filled with 700 µl of TRIzol^TM^ (Thermo Fisher Scientific, Waltham, MA) and spun in a bullet blender (Next Advance, Troy, NY) for 5 min at full speed. The lysate was retrieved, transferred to a new tube, and left for 5 min at room temperature. It was then mixed with 140 µl of chloroform, left for 3 min at room temperature, and centrifuge at 12.000 g for 15 minutes, at 4 °C. Next, the supernatant was transferred to a new tube and mixed with 1.5 volume of 100% ethanol. The mix was loaded on an RNA Purification Column and centrifuged at 8.000 g for 15 s at room temperature. Then, 10 µl of DNAse and 30 µl of RDD buffer (Qiagen, Germany) were added on the column and at room temperature for 15 min. The column was then washed one time with 700 µl of RWT buffer (8.000, 15s, room temperature), 2 times with 500 µl of RPE buffer (8.000, 15s, room temperature) and spined another time for 1 min at full speed to dry the membrane. The column was placed in a new 1.5 mL collection tube and 30 of RNase-free water was loaded on the membrane. Finally, the RNA was eluted by centrifugation for 1 min at 8.000 g. RNA concentrations were measured by spectrophotometry (Nanodrop One, Thermo Fisher Scientific, Waltham, MA). Samples were then kept at −80 °C until use.

### RT-qPCR

To generate cDNA, 50 ng of the total mRNA was reversed transcribed with a qScript cDNA synthesis kit (QuantaBio, Beverly, Massachusetts) following manufacturer’s instructions. Real-time quantitative polymerase chain reaction (RT-qPCR) was performed in 10 μL reaction containing 2 μL of cDNA (1/10 dilution), 5 μL of PowerUp SYBR Green Master Mix (Applied Biosystems, Foster City, California, United-States), 2 μL of a mix of forward and reverse primers (10 μM) and 1 μL of H2O. The cycling conditions were 50 °C for 2 min, 95 °C for 10 min, and then 40 cycles at 95 °C for 15 s and 60 °C for 1 min. The following primers were used:

*B2m*: F: TGCTACGTAACACAGTTCCACC; R: TCTGCAGGCGTATGTATCAGTC

*Dbh*: F: CCGAAATGCCAAAATTGTCA; R: GGACCCCTGCCTGTATTTTGT

*Th*: F: TGCAGCCCTACCAAGATCAAAC ; R: CGCTGGATACGAGAGGCATAGTT

*Adra1a*: F: TGCAGCCCTACCAAGATCAAAC ; R: CGCTGGATACGAGAGGCATAGTT

*Adra2a*: F: TGCAGCCCTACCAAGATCAAAC ; R: CGCTGGATACGAGAGGCATAGTT

*Adrb2*: F: TGCAGCCCTACCAAGATCAAAC ; R: CGCTGGATACGAGAGGCATAGTT

*Crhr1*: F: GGAACCTCATCTCGGCTTTCA ; R: GTTACGTGGAAGTAGTTGTAGGC

Data were normalized to *B2m* (from the same animal), and fold changes were calculated using the 2^-ΔΔCt^ method [98].

### Statistics and Data Analysis

All summary data are expressed as mean ± SEM. Statistical significance was taken as *p < 0.05, **p < 0.01, ***p < 0.001, as determined by the Student’s t-test (paired and unpaired): One-Way Analysis of Variance (ANOVA) or One-Way Repeated Measures ANOVA, followed by Dunnett’s or Bonferroni post hoc tests as appropriate. In cases where data failed the D’Agostino and Pearson omnibus normality test, non-parametric analyses were used. Statistical analyses were performed in GraphPad Prism 10. Immunohistochemistry staining of cFos was analyzed using ImageJ. In ImageJ, an ROI was created around the locus coeruleus and copied onto the channel that had the cFos staining. After, a threshold for the cFos intensity was created. ImageJ was then used to make a binary image of the cFos and ROI, section out overlapping cFos signal, and then count the cFos only located within the ROI.

## Acknowledgements

We thank the other members of the Al-Hasani and McCall labs for helpful feedback on this project. Special thanks to Patricia Jensen for the *Dbh^Cre^* mice. This work was financially supported by the National Institutes of Health (R01NS117899, J.G.M.; R01NS135401, R.A., J.G.M.), the National Science Foundation (DGE-2139839, C.R.V.), a Collaboration Support initiative for Translational Anesthesiology Research (COSTAR) award from the Department of Anesthesiology at Washington University School of Medicine (J.G.M.), and IDDRC@WUSTL (NICHD P50HD103525, S.E.M.). We would like to acknowledge biorender.com for figure cartoons, the Washington University School of Medicine Hope Center for Neurological Disorders viral vector core, and the Osage Nation, Missouria, Illinois Confederacy, and many other tribes as the ancestral, traditional, and contemporary custodians of the land where this work was conducted.

## Author contributions

C.R.V. and J.G.M. conceived the project and designed the detailed experimental protocols. C.R.V., L.J.B., C.-C.K., S.A.C., and A.N.H. performed the mouse experiments. C.R.V., L.J.B., C.-C.K., S.A.C., A.N.H., S.E.M., and J.G.M performed the investigation and analyzed the data. C.R.V., L.J.B., and J.G.M. wrote the paper. C.R.V., L.J.B., C.-C.K., S.E.M., and J.G.M. edited the paper. C.R.V., R.A., S.E.M., and J.G.M. acquired funding. R.A., J.G.M. provided research supervision. J.G.M. led overall project administration. All authors discussed the results and contributed to revision of the manuscript.

## Conflict of Interest

The authors declare no conflicts of interest.

---

**Supplementary Figure 1:**
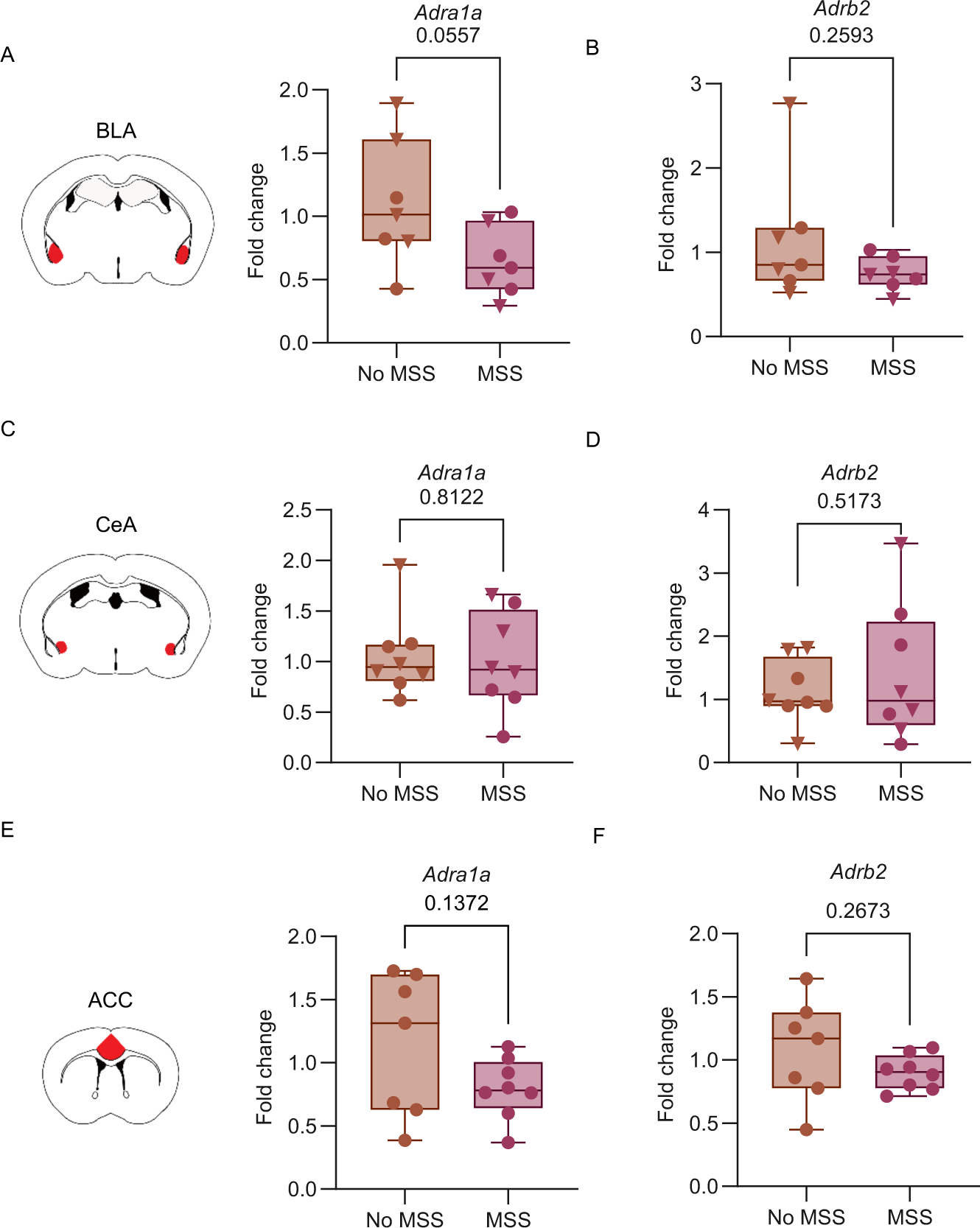
mRNA Expression in LC projection areas in MSS and No MSS mice. (**A**) In the BLA, *Adra1a* expression displayed a downward trend in MSS animals compared to animals with No MSS (unpaired t-test, t(12) = 2.118, p= 0.0557). (**B**) *Adrb2* expression in the BLA was comparable between MSS animals and No MSS animals (Mann-Whitney test, U=15, p= 0.2593). (**C**) In the CeA, *Adra1a* expression was comparable between MSS animals and no MSS (unpaired t-test, t(14) = 0.2421, p= 0.8122). (**D**) *Adrb2* expression in the CeA was comparable between MSS animals and No MSS animals (unpaired t-test, t(14) = 0.6644, p= 0.5173). (**E**) In the ACC, *Adra1a* expression was comparable between MSS animals and No MSS animals (unpaired t-test, t(13) = 1.584, p= 0.1372). (**F**) *Adrb2* expression in the ACC was comparable between MSS animals and No MSS animals (unpaired t-test, t(13) = 1.159, p=0.2673). Data are presented as box and whiskers plots, where the box represents the IQR, the line within the box indicates the median, and whiskers extend from the minimum to maximum values.

**Supplementary Figure 2:**
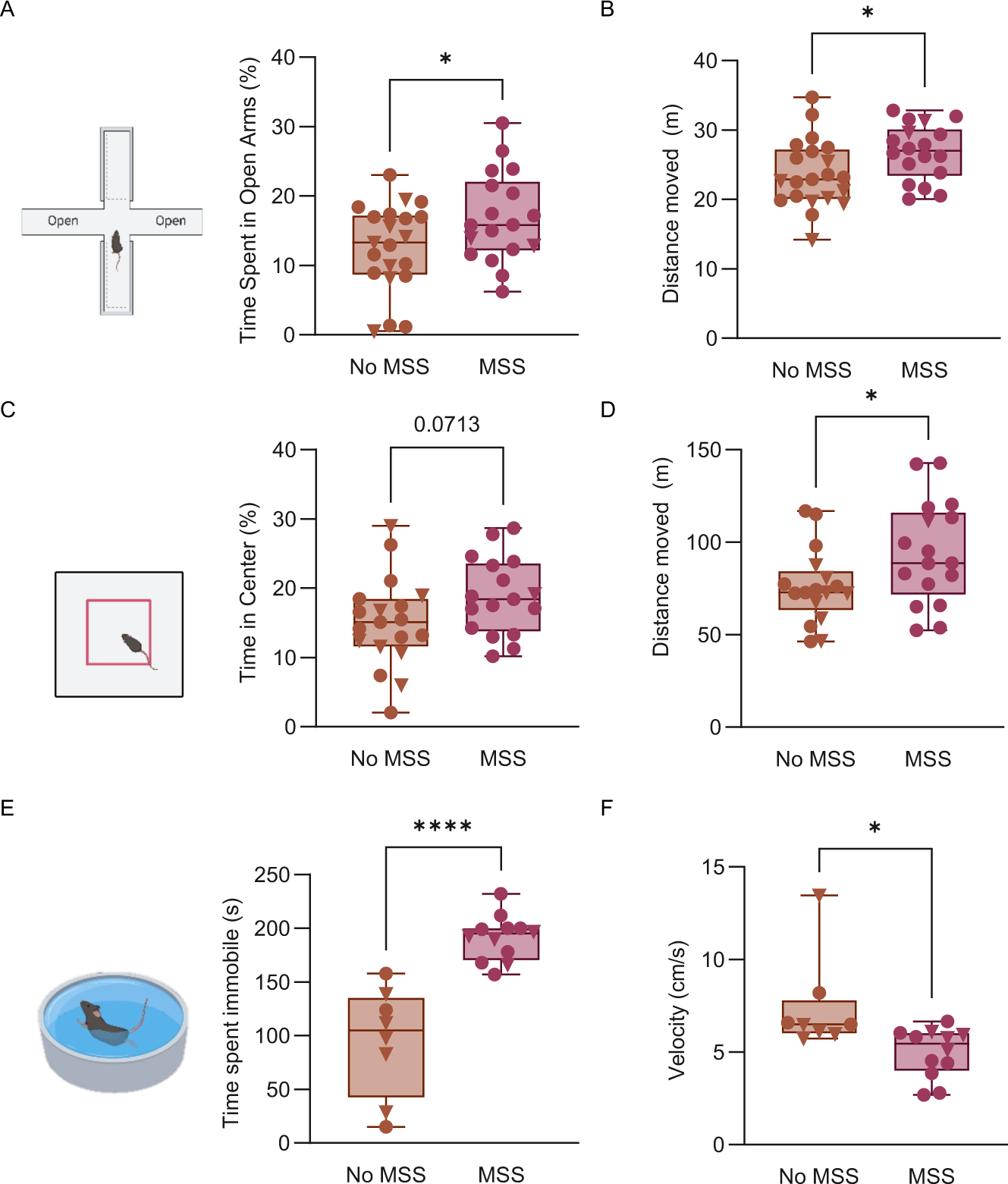
EPM, OFT, and FST data not separated by sex. (**A**) Time spent in open arms in the elevated-plus maze for MSS and No MSS groups (unpaired t-test, t(37) = 2.098, p= 0.0428). (**B**) Distance traveled in the elevated-plus maze for MSS and No MSS groups (Unpaired t-test, t(37) = 2.171, p= 0.0364). (**C**) Time spent in the center in the open field test for MSS and No MSS groups (Unpaired t-test, t(34) = 1.861). (**D**) Distance traveled in the open field test for MSS and No MSS groups (Mann-Whitney, U=84, *p=0.0376). (**E**) Immobility in the forced swim test for MSS and No MSS groups (Unpaired t-test, t(18) = 5.924, ****p<0.0001). (**F**) Velocity in the forced swim test for MSS and No MSS groups (Unpaired t-test, t(18) = 2.760, p=0.0129). Data are presented as box and whiskers plots, where the box represents the IQR, the line within the box indicates the median, and whiskers extend from the minimum to maximum values.

**Supplementary Figure 3:**
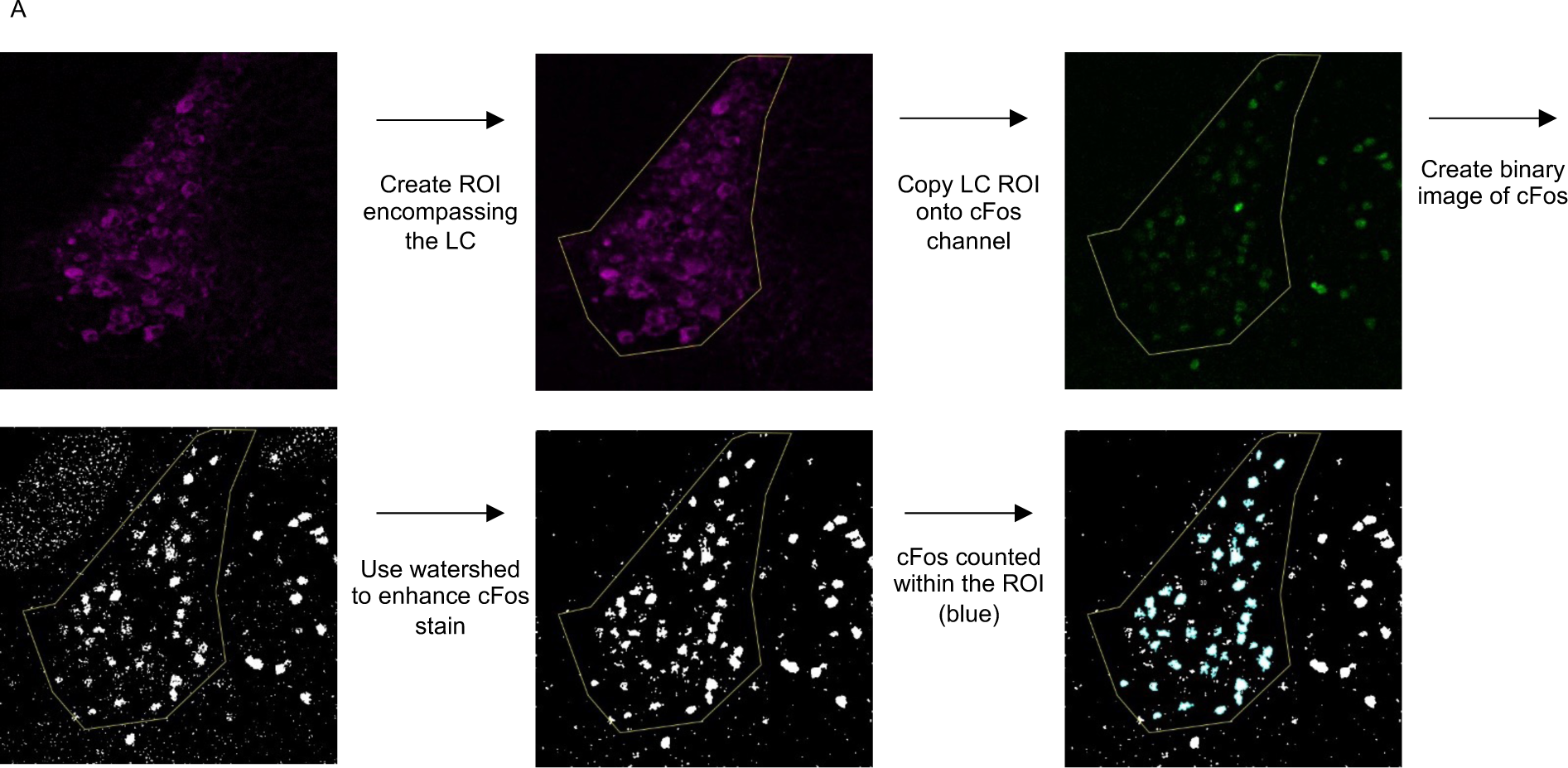
Using ImageJ to analyze cFos in the LC region. Schematic of the process to analyze cFos after immunohistochemistry staining as described in the methods.

## Notes

### Competing Interest Statement

The authors have declared no competing interest.

